# DeepGS: Predicting phenotypes from genotypes using Deep Learning

**DOI:** 10.1101/241414

**Authors:** Wenlong Ma, Zhixu Qiu, Jie Song, Qian Cheng, Chuang Ma

## Abstract

**Motivation:** Genomic selection (GS) is a new breeding strategy by which the phenotypes of quantitative traits are usually predicted based on genome-wide markers of genotypes using conventional statistical models. However, the GS prediction models typically make strong assumptions and perform linear regression analysis, limiting their accuracies since they do not capture the complex, non-linear relationships within genotypes, and between genotypes and phenotypes.

**Results:** We present a deep learning method, named DeepGS, to predict phenotypes from genotypes. Using a deep convolutional neural network, DeepGS uses hidden variables that jointly represent features in genotypic markers when making predictions; it also employs convolution, sampling and dropout strategies to reduce the complexity of high-dimensional marker data. We used a large GS dataset to train DeepGS and compare its performance with other methods. In terms of mean normalized discounted cumulative gain value, DeepGS achieves an increase of 27.70%~246.34% over a conventional neural network in selecting top-ranked 1% individuals with high phenotypic values for the eight tested traits. Additionally, compared with the widely used method RR-BLUP, DeepGS still yields a relative improvement ranging from 1.44% to 65.24%. Through extensive simulation experiments, we also demonstrated the effectiveness and robustness of DeepGS for the absent of outlier individuals and subsets of genotypic markers. Finally, we illustrated the complementarity of DeepGS and RR-BLUP with an ensemble learning approach for further improving prediction performance.

**Availability:** DeepGS is provided as an open source R package available at https://github.com/cma2015/DeepGS.

## 1 Introduction

Genomic selection (GS), originally proposed by Meuwissen *et al*. (2001) for animal breeding, is regarded as a promising breeding paradigm to better predict the plant or crop phenotypes of polygenic traits by using genome-wide markers (Bhat, *et al*., 2016; Desta and Ortiz, 2014; Jonas and De Koning, 2013; Poland and Rutkoski, 2016). Unlike both phenotypic and traditional marker-based selection, GS has the inherent advantages of predicting phenotypic trait values of individuals before planting, of estimating the breeding values of individuals before crosses are made, and, notably, of reducing the time length of the breeding cycle (Desta and Ortiz, 2014; Jannink, *et al*., 2010; Jonas and De Koning, 2013; Yu, *et al*., 2016). Recently, several GS projects have been launched for crop species, namely wheat, maize, rice and cassava (Guzman, *et al*., 2016; Marulanda, *et al*., 2016; Poland and Rutkoski, 2016; Spindel, *et al*., 2015). However, the application of GS in the field of practical crop breeding is still nascent, largely because it must overcome the requirement of robust approaches for making accurate predictions in high-dimensional datasets, where the number of genotypic markers (*p*) is much larger than the population size (*n*) (*p* >> n) (Crossa, *et al*., 2017; Desta and Ortiz, 2014; Jannink, *et al*., 2010; Schmidt, *et al*., 2016).

Various statistical models have been developed for GS, including BLUP (best linear unbiased prediction)-based algorithms, such as the ridge regression BLUP (RR-BLUP) (Endelman, 2011) and the genomic relationship BLUP (GBLUP) (VanRaden, 2008), and Bayesian-based algorithms, such as Bayes A, Bayes B, Bayes Cπ and Bayes LASSO (De Los Campos, *et al*., 2009; Meuwissen, *et al*., 2001). However, among the different statistical models, not much variation in prediction accuracy was frequently observed (Roorkiwal, *et al*., 2016; Varshney, 2016). In addition, the statistical models typically make strong assumptions and perform linear regression analysis. A representative example is the commonly used RR-BLUP model, which assumes that all the marker effects are normally distributed with a small but non-zero variance, and predicts phenotypes from a linear function of genotypic markers (Xu and Crouch, 2008). As a result, the GS models based on traditional statistical methods not only have to face the statistical challenges related to the high dimensionality of marker data, but also have difficulty capturing complex relationships within genotypes (e.g., multicolinearity among markers), and between genotypes and phenotypes (e.g., genotype-by-environment-by-trait interaction) (Crossa, *et al*., 2017; Van Eeuwijk, *et al*., 2010). Therefore, novel methods are urgently needed to augment GS and its potential in plant breeding.

Deep learning (DL) is a recently developed machine-learning technique that provides good prediction capability with many advanced features, one of which is the deep multi-layered neural network architecture. In a deep multi-layered neural network, a large number of neurons are used to capture complex, nonlinear relationships in big data (large datasets) (LeCun, *et al*., 2015). DL has proven capable of improved prediction performance over traditional models for speech recognition, image identification and natural language processing (LeCun, *et al*., 2015). Most recently, however, DL has drawn the attention of systems biologists, who have successfully applied it to several prediction problems: the inference of gene expression (Chen, *et al*., 2016; Singh, *et al*., 2016), the functional annotation of genetic variants (Quang, *et al*., 2015; Quang and Xie, 2016; Xiong, *et al*., 2015; Zhou and Troyanskaya, 2015), the recognition of protein folds (Jo, *et al*., 2015; Wang, *et al*., 2016) and the prediction of genome accessibility (Kelley, *et al*., 2016), of enhancers (Kim, *et al*., 2016; Liu, *et al*., 2016), and of DNA- and RNA-binding proteins (Alipanahi, *et al*., 2015; Zeng, *et al*., 2016; Zhang, *et al*., 2016). These successful applications in the fields of computational biology and systems biology have demonstrated that DL has a powerful capability of learning complex relationships from biological data (Angermueller, *et al*., 2016; Min, *et al*., 2017). However, to the best of our knowledge, the application of DL in the field of GS has not yet been investigated.

In this study, we present a DL method, named DeepGS, to predict phenotypes from genotypes by using a deep convolutional neural network (CNN). Unlike the conventional statistical models, DeepGS can automatically “learn” complex relationships between genotypes and phenotypes from the training dataset, without pre-defined rules (e.g., normal distribution, non-zero variance) for various variables in the neural network. In order to avoid overfitting of CNN, DeepGS also takes the advantages of DL technologies to reduce the complexity of high-dimensional marker data through dimensionality reduction using convolution, sampling and dropout strategies. We used a large GS data of wheat (2,000 individuals × 33,709 markers; eight phenotypic traits) from CIMMYT (International Maize and Wheat Improvement Center), to train DeepGS and compare its performance to those of other models. Cross-validation experimental results showed that the DeepGS outperformed a conventional feed-forward neural network in the prediction of phenotypic values for all eight tested traits. DeepGS also had superiority over the widely used GS method RR-BLUP in selecting individuals with high phenotypic values. Further simulation experiments indicated that DeepGS still had the advantage over RR-BLUP in selecting individuals with high phenotypic values, even for the absent of outlier individuals and subsets of genotypic markers. We also proposed an ensemble learning approach to linearly combine the predictions of DeepGS and RR-BLUP for further improving the prediction performance. These results suggest that DeepGS can be used as a supplementary to RR-BLUP for selecting individuals with high phenotypic values. DeepGS has been implemented as an open source R package now available for public use (https://github.com/cma2015/DeepGS).

## 2 Methods

### 2.1 GS dataset

The GS dataset used in this study was obtained from the wheat gene bank of CIMMYT, which consists of 2,000 Iranian bread wheat (*Triticum aestivum*) landrace accessions genotyped with 33,709 DArT (Diversity Array Technology). For the DArT markers, an allele was encoded by either 1 or 0, to indicate its presence or absence, respectively. Each of these accessions was phenotyped for eight traits: grain length (GL), grain width (GW), grain hardness (GH), thousand-kernel weight (TKW), test weight (TW), sodium dodecyl sulphate-sedimentation (SDS), grain protein (GP) and plant height (PHT). More information about this GS dataset was presented in a recently published paper (Crossa, *et al*., 2016). The complete genotypic and standardized phenotypic datasets can be obtained from http://genomics.cimmyt.org/mexican_iranian/traverse/iranian/standarizedData_univariate.RData.

### 2.2 10-fold cross-validation

Cross-validation has been used to evaluate the prediction performance of GS models (Crossa, *et al*., 2016; Gianola and Schon, 2016; Qiu, *et al*., 2016; Resende, *et al*., 2012). In this study, a 10-fold cross-validation has been used, in which individuals in the whole GS dataset were first randomly partitioned into 10 groups with approximately equal size. The GS model was trained and validated using genotypic and phenotypic data of individuals from nine groups (90% individuals for the training set; 10% individuals for the validation set). The trained GS model was subsequently applied to predict phenotypic trait values of individuals from the remaining group (testing set) using only genotypic data. This process was repeated 10 times until each group was used once for testing; the predicted phenotypic trait values were finally combined for performance evaluation.

The prediction performance of each GS model for selecting individuals with high phenotypic values is assessed by the measure: the mean normalized discounted cumulative gain value (MNV) (Blondel, *et al*., 2015). Given *n* individuals, the predicted and observed phenotypic values form an *n* × 2 matrix of score pairs (*X, Y*). The MNV for selecting the top-ranked *k* individuals can be calculated in an iterative manner:

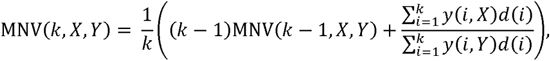
where, *d*(*i*) = 1/(log_2_ *i* + 1) is a monotonically decreasing discount function at position *i*; *y*(*i*,*Y*) is the *i_th_* value of observed phenotypic values *Y* sorted in descending order, here *y*(1,*Y*) ≥ *y*(*2*,*Y*) ≥ … ≥ *y*(*n*,*Y*); *y*(*i*,*X*) is the corresponding value of *Y* in the score pairs (*X*, *Y*) for the *i_th_* value of predicted scores *X* sorted in descending order. Thus, MNV has a range of 0 to 1 when all the observed phenotypic values are larger than zero; a higher MNV(*k, X, Y*) indicates a better performance of the GS model to select the top-ranked *k* individuals with high phenotypic values.

### 2.3 Ridge regression-based linear unbiased prediction (RR-BLUP)

RR-BLUP is one of the most extensively used and robust regression models for GS (Bhering, *et al*., 2015; Huang, *et al*., 2016; Wimmer, *et al*., 2013). Given the genotype matrix *Z* (*n* × *p*; *n* individuals, *p* markers) and the corresponding phenotype vector *Y* (*n* × 1), the GS model is built using the standard linear regression formula:

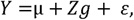
where, *μ* is the mean of phenotype vector *Y*, *g*(*p* × 1) is a vector of marker effects, and *ε* (*n* × 1) is the vector of random residual effects. The ridge regress algorithm is used to simultaneously estimate the effects of all genotypic markers, under the assumption that marker effects in *g*(*p* × 1) follow a normal distribution norm 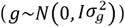 with a small but non-zero variance 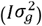 (Desta and Ortiz, 2014; Endelman, 2011; Riedelsheimer, *et al*., 2012; Whittaker, *et al*., 2000). *I* is the identity matrix, 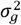 is the variance of *g*. The RR-BLUP model was implemented using the function “mixed.solve” in the R package “rrBLUP”

### 2.4 DeepGS model

The DeepGS model was built using the DL technique-deep convolutional neural network (CNN) with an 8-32-1-architecture; this included one input layer, one convolutional layer (eight neurons), one sampling layer, three dropout layers, two fully-connected layers (32 and one neurons) and one output layer (Fig. 1). The input layer receives the genotypic markers of a given individual in the 1 × *p* matrix, where *p* is the number of genotypic markers. The first convolutional layer filters the input matrix with eight kernels that are each 1 × 18 in size with a stride size of 1 × 1, followed by a 1 × 4 max-pooling layer with a stride size of 1 × 4. The output of the max-pooling layer is passed to a dropout layer with a rate of 0.2 for reducing overfitting (Srivastava, *et al*., 2014). The first fully-connected layer with 32 neurons is used after the dropout layer to join together the convolutional characters with a dropout rate of 0.1. A nonlinearity active function - rectified linear unit (ReLU), is applied in the convolutional and first fully connected layers. The output of the first fully-connected layer is then fed to the second fully-connected layer with one neural and a dropout rate of 0.05. By using a linear regression model, the output of the second fully-connected layer is finally connected to the output layer which presents the predicted phenotypic value of the analyzed individual.

**Fig. 1.**
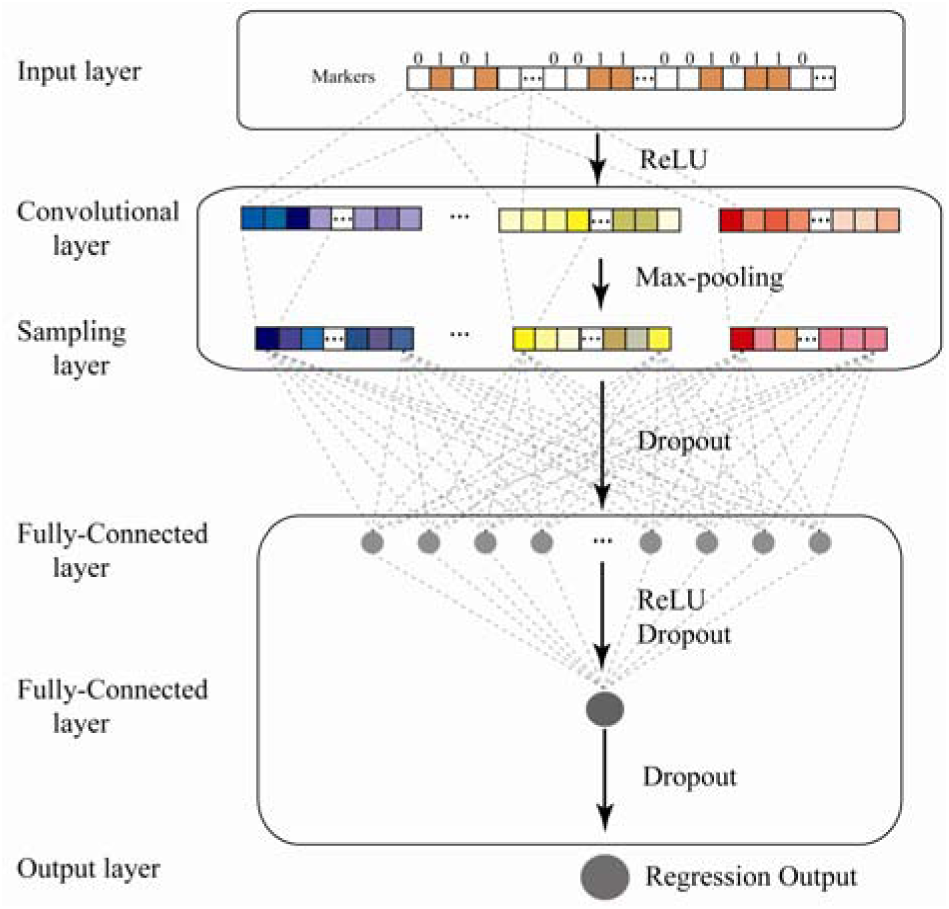
The DeepGS model is a deep convolutional neural network that has an 8-32-1 architecture. “Dropout” denotes the dropout layer. ‘ReLU’ indicates the rectified linear unit. (https://cran.r-project.org/web/packages/rrBLUP).

For each fold of cross-validation, the DeepGS was trained on the training set and validated on the validation set. Parameters in the DeepGS were optimized with the back propagation algorithm (Rumelhart, *et al*., 1986), by setting the number of epochs to 6,000, the learning rate to 0.01, the momentum to 0.5, and the *wd* to 0.00001. The loss function we minimized is the mean absolute error (Mae) index:

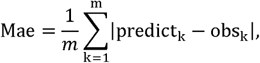
where, *m* denotes the number of individuals in the training dataset, and *predict_k_* and *obs_k_* represent the predicted and observed phenotypic values of the *k_th_* individual, respectively.

DeepGS was implemented using the graphics-processing-unit (GPU)-based DL framework MXNet (version 0.7.0; https://github.com/dmlc/mxnet); it was run on a GPU server that was equipped with four NVIDIA GeForce TITAN-XGPUs, each of which has 12GB of memory and 3072 CUDA (Compute Unified Device Architecture) cores.

### 2.5 An integrated GS model linearly combining RR-BLUP and DeepGS

An integrated GS model (*I*) was constructed using the ensemble learning approach by linearly combining the predictions of DeepGS (*D*) and RR-BLUP (*R*), using the formula:

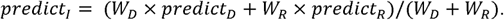

For each fold of 10-cross fold procedure, parameters (*W_D_* and *W_R_*) were optimized on the corresponding validation dataset using the particle swarm optimization (PSO) algorithm, which was developed by inspiring from the social behavior of bird flocking or fish schooling (Kennedy and Eberhart, 1995). PSO has the capability of parallel searching on very large spaces of candidate solutions, without making assumptions about the problem being optimized. Details of the parameter optimization using the PSO algorithm are given in **Supplementary Information**.

### 2.6 Statistical analysis in this study

The Pearson’s correlation coefficient (PCC) and corresponding significance level (*p*-value) were calculated with the function “cor.test” in R programming language (https://www.r-project.org). The significance level of the difference between paired samples was examined using the student’s t-test with R function “t.test”.

## 3 Results

### 3.1 DeepGS outperforms a conventional neural network and the random selection

To perform the regression-based GS using neural network algorithms, we were interested in whether or not the DL-based neural network model (DeepGS) was more powerful than the conventional neural network model. To address this task, a three-layer, fully-connected, feed-forward neural network (FNN) was built using the matlab function “feedforwardnet”, in which there was also an 8-32-1 architecture (i.e. eight nodes in the first hidden layer, 32 nodes in the second hidden layer, and one node in the output layer). In FNN, nodes in one layer were fully connected to all nodes in the next layer. The 10-fold cross-validation was performed to evaluate the performance of DeepGS and FNN for predicting phenotypic values of the eight tested traits using 33,709 DArT markers.

For the trait of grain length (GL), PCC analysis revealed that these two GS models had predicted phenotypic values that were significantly correlated with the observed phenotypic values (student’s t-test; *p*-value < 1.00E-69) (Fig. 2A-2B). However, the DeepGS had a markedly higher PCC value (0.745) than did the FNN (0.378) (Fig. 2A-2B). Correspondingly, the predictions of DeepGS had a significantly lower absolute error compared with that for the FNN (paired samples t-test; *p*-value <2.00E-66) (Fig. 2C).

**Fig. 2.**
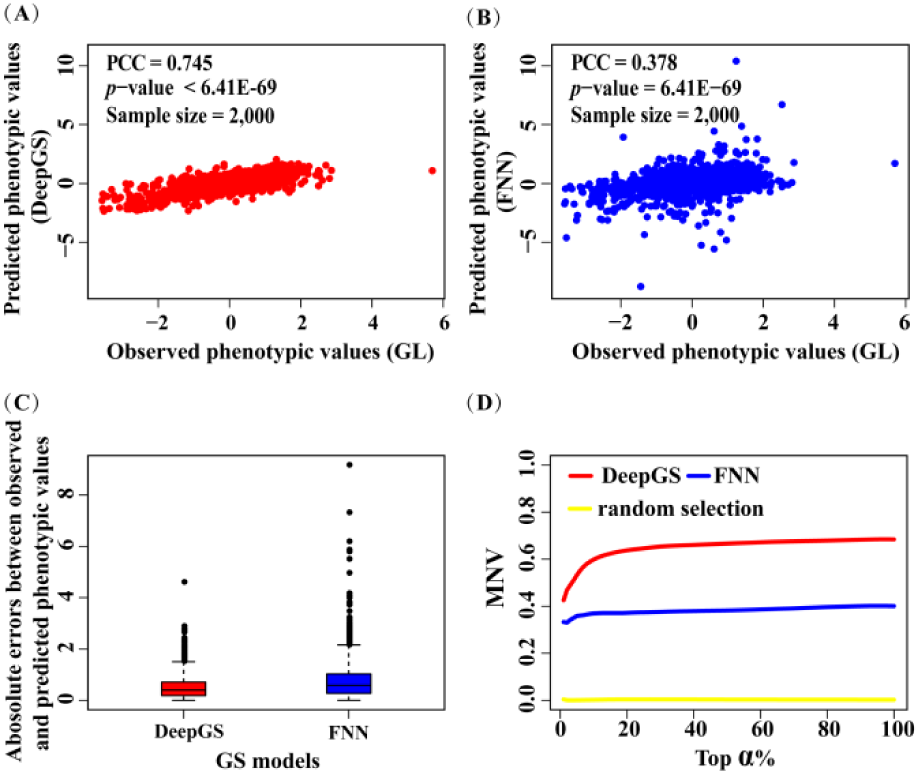
Performance of DeepGS and FNN for predicting the grain length of wheat using 33,709 DArT markers. (**A**) Dot-plot comparison of observed phenotypic values with predicted phenotypic values of the DeepGS, and (**B**) the FNN. (**C**) Boxplot of the absolute errors between the observed and predicted phenotypic values. (**D**) MNV curves for DeepGS, FNN, and random selection with top-ranked *α* increasing from 1% to 100%.

The MNV was further used to evaluate the performance of the DeepGS and FNN GS models for selecting individuals with high grain length. The MNV of the DeepGS model (0.43~0.68) was significantly higher than that of the FNN-based GS model (0.33~0.40) (paired samples t-test; *p*-value < 7.50E-91), with top-ranked *α* increasing from 1% to 100% (Fig. 2D). Both DeepGS and FNN had markedly higher MNVs than those generated from random selection (0~0.0040). In the random selection experiment, individuals were randomly ranked from 1 to 2,000, and this process was repeated 100 times, which generated 100 MNVs for each given *α*. The mean of these 100 MNVs was used to quantify the final performance of the random selection for the given *α*. For the other seven traits under study, we also observed that the performance followed the order of: DeepGS > FNN > random selection (**Supplementary Fig. S1**). At the top-ranked level of *α* = 1%, the MNV improvement of DeepGS over FNN could be as high as 74.37%, 58.98% (*α* = 1%), 89.10% (*α* = 2%), 62.49% (*α* = 18%), 86.92% (*α* = 15%), 158.68% (*α* = 3%), 150.92% (*α* = 8%), and 445.71% (*α* = 8%) for GL, GW, GH, TKW, TW, SDS, GP, and PHT, respectively.

Taken together, these results showed that DeepGS outperforms both the FNN and the random selection for predicting phenotypic values of the eight tested traits.

### 3.2 DeepGS outperforms RR-BLUP for selecting individuals with high phenotypic values

For each of the eight traits under study, we performed 10-fold cross-validation to evaluate the performance of RR-BLUP and DeepGS for selecting individuals with high phenotypic values. Paired samples t-test analysis showed that, when *α* ranged from 1% to 100%, the MNVs of DeepGS model were significantly higher than those of the RR-BLUP for all tested traits except PHT (Table 1; Fig. 3A). The relative improvement of DeepGS over RR-BLUP in MNVs could be as high as 19.94% (*α* = 1%), 23.72% (*α* = 1%), 3.60% (*α* = 5%), 36.11% (*α* = 1%), 37.34% (*α* = 1%), 6.15% (*α* = 2%), 15.70% (*α* = 1%), and 65.24% (*α* = 1%) for GL, GW, GH, TKW, TW, SDS, GP and PHT, respectively (Table 1). These results indicate that DeepGS outperformed RR-BLUP, especially for selecting individuals with extremely high phenotypic values of the eight tested traits.

**Table 1.** Prediction performance of DeepGS and RR-BLUP for eight tested traits. “integrated” indicates the integrated GS model. “*p*-value” represents the significance level of MNV improvement.

| Trait | MNVs |  |  | DeepGS vs RR-BLUP |  | integrated vs RR-BLUP |  |
| --- | --- | --- | --- | --- | --- | --- | --- |
|  | RR-BLUP | DeepGS | integrated | MNV improvement (%)<br>(median) | *p*-value | MNV improvement (%)<br>(median) | *p*-value |
| GL | 0.35~0.67 | 0.43~0.68 | 0.39~0.69 | 1.06~19.94 (3.39) | 2.15E-59 | 0.38~9.08 (3.52) | 3.84E-73 |
| GW | 0.62~0.77 | 0.75~0.80 | 0.71~0.79 | 1.71~23.72 (1.97) | 3.18E-21 | 1.65~14.68 (1.88) | 7.52E-32 |
| GH | 0.70~0.76 | 0.70~0.78 | 0.71~0.78 | 0.10~3.60 (1.00) | 3.48E-25 | 1.39~3.49 (1.94) | 1.97E-63 |
| TKW | 0.47~0.66 | 0.64~0.68 | 0.64~0.69 | 2.62~36.11 (3.13) | 7.48E-23 | 3.58~36.51 (3.91) | 8.05E-29 |
| TW | 0.23~0.61 | 0.32~0.62 | 0.31~0.62 | 1.44~37.34 (2.24) | 1.55E-31 | 2.19~33.30 (2.95) | 2.08E-45 |
| SDS | 0.56~0.74 | 0.56~0.78 | 0.57~0.76 | -1.36~6.15 (-0.20) | 9.71E-04 | 1.09~3.77 (1.75) | 2.07E-45 |
| GP | 0.27~0.50 | 0.31~0.50 | 0.35~0.51 | 0.21~15.70 (1.04) | 1.29E-17 | 3.25~29.48 (3.89) | 1.27E-44 |
| PHT | 0.26~0.31 | 0.25~0.42 | 0.27~0.43 | -10.68~65.24 (-8.75) | 1.00E-00 | -1.53~67.04 (-0.35) | 3.68E-03 |

**Fig. 3.**
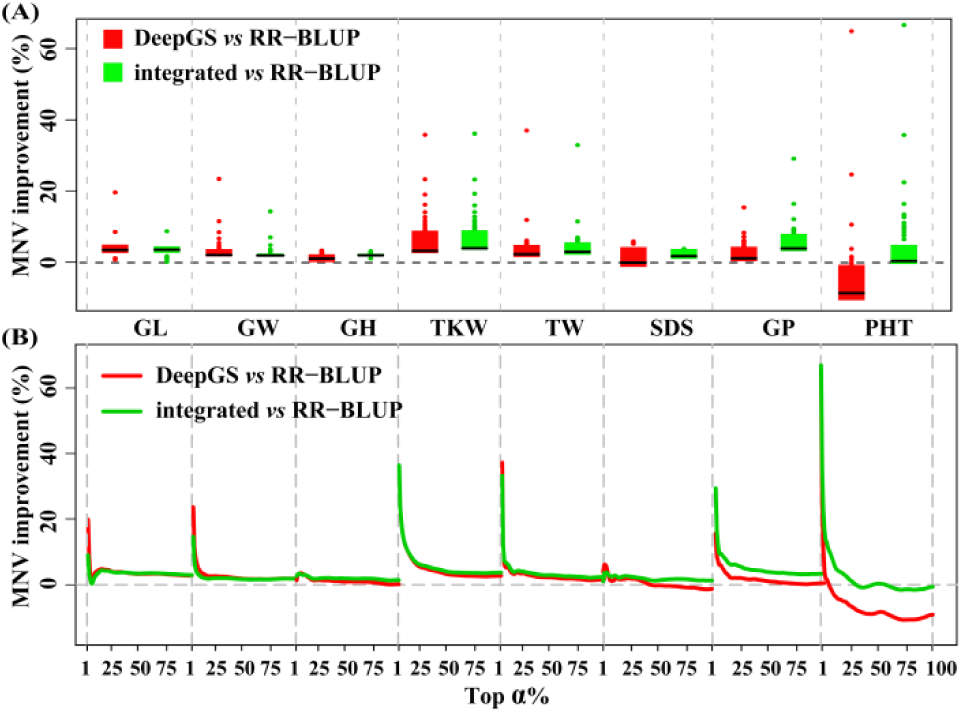
The MNV improvement of DeepGS and the integrated GS models over RR-BLUP for eight tested traits. (**A**) Boxplot of the MNV improvement with top-ranked *α* increasing from 1% to 100%. (**B**) The curves of MNV improvement with top-ranked *α* increasing from 1% to 100%.

Considering that DeepGS and RR-BLUP used different algorithms to build regression-based GS models, we suspected that these two approaches may capture different aspects of the relationships between genotypes and phenotypes. Thus, the combination of predictions of DeepGS and RR-BLUP may contribute to a better performance. As expected, in terms of MNV, the integrated GS model also gained significant higher performance than RR-BLUP for all tested traits when top-ranked *α* ranged from 1% to 100% by paired samples t-test (Table 1; Fig. 3B). Obviously, the integrated GS model substantially improved the prediction performance over RR-BLUP and DeepGS for GP and PHT (Fig. 3B). Compared with RR-BLUP, DeepGS improved the MNVs by 0.21%~15.70% for GP and of -10.68%~65.24% for PHT; while the integrated GS model improved the MNVs by 3.25%~29.48% for GP and -1.53%~67.04% for PHT (Table 1).

These results indicated that the DeepGS can be used as a supplementary to the RR-BLUP model in selecting individuals with high phenotypic values for all of the eight tested traits.

### 3.3 Outlier individuals and their effects on prediction performance

An outlier individual is one with an extremely high or low phenotypic value for a particular trait under study. These outlier individuals are valuable for breeding programs and for identifying trait-related genes in the bulked sample analysis (Zou, *et al*., 2016). We were interested in how the respective performance of DeepGS and RR-BLUP models might be affected by outlier individuals. For each of the eight traits, the outlier individuals were defined as above 75% quartile (Q3) plus 1.5 times the interquartile range (IQR = Q3 – Q1) and below 25% quartile (Q1) minus 1.5 times IQR of phenotypic values. We removed 50, 22, 40, 19, and 65 outlying individuals for GL, GW, TW, GP and PHT, respectively (**Supplementary Fig. S2A**). The remaining individuals of these five traits were used to evaluate the performance of RR-BLUP, DeepGS, and the integrated GS model using the 10-fold cross validation approach.

We observed that RR-BLUP and DeepGS are differentially sensitive to outlier individuals. The removal of outlier individuals improved the MNVs of RR-BLUP at different levels of *α* for GL (1% ≤ *α* ≤ 100%), GW (1% ≤ *α* ≤ 2%), TW (1% ≤ *α* ≤ 100%), GP (1% ≤ *α* ≤ 36%) except PHT, and while for DeepGS, higher performances were evident for GL (1% ≤ *α* ≤ 100%), TW (1% ≤ *α* ≤ 100%), and GP (1% ≤ *α* ≤ 100%) except GW and PHT (**Supplementary Fig. S2B**). However, after the removal of outlier individuals, DeepGS still yielded a higher prediction performance than it did by RR-BLUP for all tested five traits at different levels of *α*: GL (1% ≤ *α* ≤ 32%), GW (1% ≤ *α* ≤ 100%), TW (1% ≤ *α* ≤ 100%), GP (1% ≤ *α* ≤ 100%), and PHT (1% ≤ *α* ≤ 15%) (Table 2; Fig. 4). The corresponding MNV improvement could be as high as 9.12%, 5.46%, 23.63%, 54.50%, and 199.48% at the level of *α* = 1% (**Supplementary Fig. S2C**). As expected, the integrated GS model always yielded a higher prediction performance than it did by RR-BLUP for all tested five traits at all possible levels of *α* (1% ≤ *α* ≤ 100%) except for PHT (1% ≤ *α* ≤ 15%) (Fig. 4; **Supplementary Fig. S2C**).

**Table 2.** Prediction performance of DeepGS and RR-BLUP for five tested traits after the removal of outlier individuals. “integrated” indicates the integrated GS model. “*p*-value” represents the significance level of MNV improvement.

| Trait | MNVs |  |  | DeepGS vs RR-BLUP |  | integrated vs RR-BLUP |  |
| --- | --- | --- | --- | --- | --- | --- | --- |
|  | RR-BLUP | DeepGS | integrated | MNV improvement (%)<br>(median) | *p*-value | MNV improvement (%)<br>(median) | *p*-value |
| GL | 0.53~0.69 | 0.58~0.70 | 0.59~0.70 | -0.30~9.12(0.39) | 5.24E-07 | 1.07~11.49 (1.51) | 1.18E-26 |
| GW | 0.66~0.75 | 0.70~0.76 | 0.69~0.76 | 0.01~5.46(0.46) | 2.64E-14 | 0.81~4.50 (1.24) | 1.05E-51 |
| TW | 0.36~0.63 | 0.44~0.64 | 0.44~0.65 | 0.97~23.63(2.63) | 2.00E-29 | 2.11~24.10 (3.38) | 2.18E-41 |
| GP | 0.31~0.49 | 0.46~0.50 | 0.44~0.51 | 2.32~54.50(3.39) | 1.87E-15 | 3.50~44.81 (4.20) | 5.25E-24 |
| PHT | 0.10~0.28 | 0.23~0.30 | 0.23~0.27 | -5.12~199.48(-4.13) | 7.52E-01 | -2.30~153.26 (-1.15) | 6.77E-02 |

**Fig. 4.**
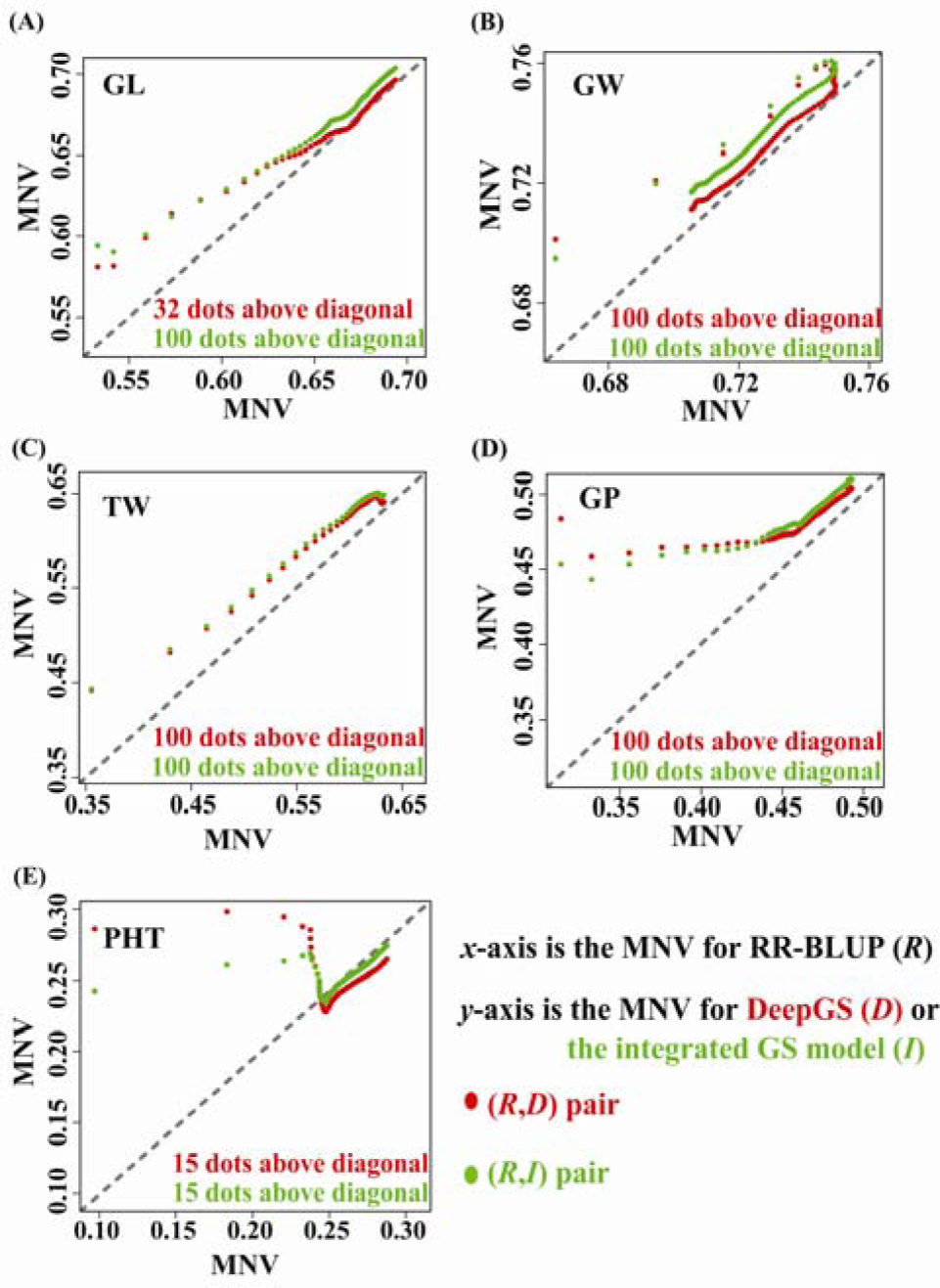
The MNVs of DeepGS and the integrated GS model compared with those of RR-BLUP for five tested traits after the removal of outlier individuals. Each point in red (or in green) represents a pair of MNVs from DeepGS and RR-BLUP (or from the integrated GS model and RR-BLUP) at a top-ranked level of *α* ranging from 1% to 100%. A dot in red (or in green) above diagonal means the DeepGS (or the integrated GS model) achieved a higher MNV compared with RR-BLUP. There were 100 dots above diagonal for DeepGS and the integrated GS model for all tested traits, with the exception of GL (32 red dots) and PHT (15 red dots and 15 green dots).

These results indicate that, even after omitting the outlier individuals, DeepGS and the integrated GS model outperforms RR-BLUP in selecting individuals with high phenotypic values for all tested traits.

### 3.4 Marker number effect on prediction performance

Various technology platforms have been developed to generate genotypic markers with different sizes. The number of genotypic markers has been reported to have significant influences on the prediction performance of GS models (Heffner, *et al*., 2011). In this analysis, we examined the effect of marker number on prediction performance of RRBLUP, DeepGS, and the integrated GS model. For each of the eight tested traits, the 10-fold cross validation experiment was performed using a different number of randomly selected markers at 5,000, 10,000, and 20,000. This process was repeated 10 times to generate 10 MNVs of a given *α* for each marker number. Their average served as the final prediction performance of the GS models.

When 20,000 markers were used, DeepGS outperformed RR-BLUP for the eight tested traits at different levels of *α*: GL (1% ≤ *α* ≤ 100%), GW (1% ≤ *α* ≤ 28%), GH (1% ≤ *α* ≤ 45%), TKW (1% ≤ *α* ≤ 100%), TW (1% ≤ *α* ≤ 100%), SDS (1% ≤ *α* ≤ 1%), GP (1% ≤ *α* ≤ 3%), and PHT (1% ≤ *α* ≤ 3%) (Fig. 5A; **Supplementary Fig. S3A**). While for the integrated GS model, the MNV improvement over RR-BLUP could be observed for all tested traits except SDS. Interestingly, the MNV improvement reached 48.28% at top-ranked *α* = 1% for PHT (Fig. 5A; **Supplementary Fig. S3A**).

**Fig. 5.**
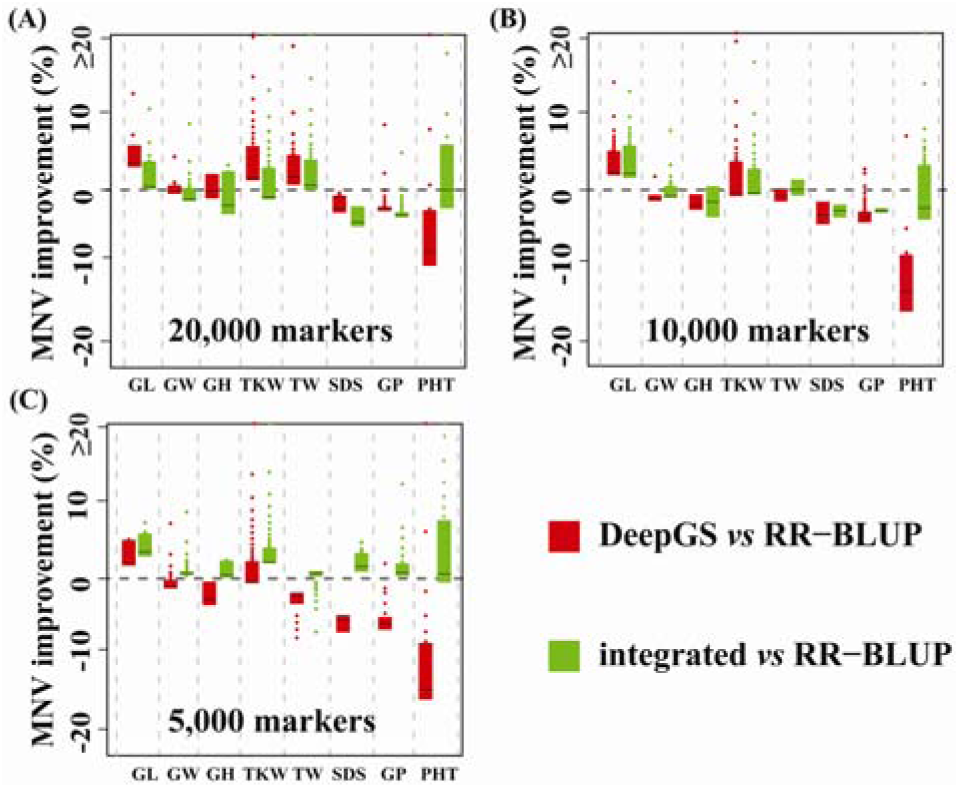
Improvement of DeepGS and the integrated GS model over RR-BLUP for eight tested traits when subsets of markers were used. Black lines represent the medians.

When the marker number decreased from 20,000 to 10,000, the MNV improvement of DeepGS over RR-BLUP was observed for GL (1% ≤ *α* ≤ 100%), GW ( 1% ≤ *α* ≤ 2%), TKW ( 1% ≤ *α* ≤ 49%), GP (1% ≤ *α* ≤ 4%), and PHT (1% ≤ *α* ≤ 1%) (Fig. 5B; **Supplementary Fig. S3B**). While for the integrated GS model, the higher performance was generated for GL (1% ≤ *α* ≤ 100%), GW (1% ≤ *α* ≤ 26%), GH (1% ≤ *α* ≤ 29%), TKW (1% ≤ *α* ≤ 48%), TW (1% ≤ *α* ≤ 79%), and PHT (1% ≤ *α* ≤ 23%) (Fig. 5B; **Supplementary Fig. S3B**).

A further decrease of the number of markers also revealed the advantage of DeepGS over RR-BLUP in selecting individuals with high phenotypic values (Fig. 5C; **Supplementary Fig. S3C**). DeepGS yielded a higher MNV than RR-BLUP for GL ( 1% ≤ *α* ≤ 100%), GW (1% ≤ *α* ≤ 17%), TKW (1% ≤ *α* ≤ 47%), GP (*α* = 1%), and PHT (1% ≤ *α* ≤ 2%). The integrated GS model further improved the prediction performance for GW, GH, TKW, SDS, and GP at all possible levels of *α* ranging from 1% to 100%, and for PHT at the levels of *α* ranging from 1% to 73% (Fig. 5C).

These results indicated that DeepGS outperforms RR-BLUP even when a subset of 33,709 markers was used and could be used as a supplementary to RR-BLUP in selecting individuals with high phenotypic values for all tested eight traits.

## 4 Discussion

GS is currently revolutionizing the applications of plant breeding, and novel prediction methods are crucial for accurately predicting phenotypes from genotypes (Desta and Ortiz, 2014; Jannink, *et al*., 2010; Jonas and De Koning, 2013). DL is a recently developed machine-learning technique, which has the capability of capturing complex relationships hidden in big data. In this study, we explored the application of DL in the field of GS. The main contributions are the following: (1) We successfully applied the DL technique to build a novel and robust GS model for predicting phenotypes from genotypes. (2) We implemented the DeepGS model as an open source R package “DeepGS”, thus providing a flexible framework to ease the application of DL techniques in GS. This R package also provides functions to calculate the MNV, and to implement the RR-BLUP model as well as the cross-validation procedure. (3) We proposed an ensemble learning approach to get a better performance through combining the predictions of DeepGS and RR-BLUP.

Nevertheless, there are several limitations to the use of DL. First, the design of appropriate network architectures is crucial to the prediction performance and requires considerable knowledge of DL and neural network. Second, the convolutional, sampling, dropout, and fully-connected layers have different sets of hyper-parameters each and thus handle different parts of the data characteristics (Angermueller, *et al*., 2016; Chen, *et al*., 2016; Min, *et al*., 2017), resulting in a challenge of interpreting and exploring biological significances. Yet, this is a general limitation of DL in the application of computational biology and bioinformatics (Min, *et al*., 2017). Recently developed network visualization systems, such as ReVACNN (https://github.com/davianlab/deepVis) and deepViz (https://github.com/bruckner/deepViz), may be helpful for providing insight into this problem. Third, extensive computational time is required to train DeepGS. Although DeepGS was implemented on a GPU server equipped with NVIDIA GeForce TITAN-X GPUs, it still required about 3.5 hours to perform the 10-fold cross-validation procedure for a single trait in the wheat GS dataset under study. To improve the running efficiency, the user could run DeepGS on a GPU-based cloud platform, e.g., Amazon Elastic Compute Cloud (Amazon EC2; https://aws.amazon.com/ec2) or Google App Engine (https://cloud.google.com/gpu).

In summary, this research work opens up a new avenue for the application of the DL technique in the field of GS. In the future, we will cooperate with population geneticists and continue to amend our DeepGS to enable it to explain the detected relationships between phenotypes and genotypes. In addition, we will cooperate with crop breeders and carry out practical applications of DeepGS in the GS-based breeding programs of wheat and other vital crops.

## Acknowledgements

Designed the experiments: CM. Performed the experiments: WM, JS, ZQ, and QC. Analyzed the data: WM, ZQ, QC and CM. Wrote the paper: CM and WM. All authors read and approved the final manuscript.

## References

Alipanahi, B., et al. Predicting the sequence specificities of DNA- and RNA-binding proteins by deep learning. Nat Biotechnol 2015;33(8):831–838.

Angermueller, C., et al. Deep learning for computational biology. Mol Syst Biol 2016;12(7):878.

Bhat, J.A., et al. Genomic selection in the era of next generation sequencing for complex traits in plant breeding. Front Genet 2016;7:221.

Bhering, L.L., et al. Comparison of methods used to identify superior individuals in genomic selection in plant breeding. Genet Mol Res 2015;14(3):10888–10896.

Blondel, M., et al. A ranking approach to genomic selection. PLoS One 2015;10(6):e0128570.

Chen, Y., et al. Gene expression inference with deep learning. Bioinformatics 2016;32(12):1832–1839.

Crossa, J., et al. Genomic prediction of gene bank wheat landraces. G3 (Bethesda) 2016;6(7):1819–1834.

Crossa, J., et al. Genomic selection in plant breeding: methods, models, and perspectives. Trends Plant Sci 2017;pii:S1360–1385(17)30184-X.

De Los Campos, G., et al. Predicting quantitative traits with regression models for dense molecular markers and pedigree. Genetics 2009;182(1):375–385.

Desta, Z.A. and Ortiz, R. Genomic selection: genome-wide prediction in plant improvement. Trends Plant Sci 2014;19(9):592–601.

Endelman, J.B. Ridge regression and other kernels for genomic selection with R package rrBLUP. Plant Genome 2011;4(3):250–255.

Gianola, D. and Schon, C.C. Cross-validation without doing cross-validation in genome-enabled prediction. G3 (Bethesda) 2016;6(10):3107–3128.

Guzman, C., et al. Wheat quality improvement at CIMMYT and the use of genomic selection on it. Appl Transl Genom 2016;11:3–8.

Heffner, E.L., Jannink, J.L. and Sorrells, M.E. Genomic selection accuracy using multifamily prediction models in a wheat breeding program. Plant Genome 2011;4(1):65–75.

Huang, M., et al. Genomic selection for wheat traits and trait stability. Theor Appl Genet 2016;129(9):1697–1710.

Jannink, J.L., Lorenz, A.J. and Iwata, H. Genomic selection in plant breeding: from theory to practice. Brief Funct Genomics 2010;9(2):166–177.

Jo, T., et al. Improving protein fold recognition by deep learning networks. Sci Rep 2015;5:17573.

Jonas, E. and De Koning, D.J. Does genomic selection have a future in plant breeding? Trends Biotechnol 2013;31(9):497–504.

Kelley, D.R., Snoek, J. and Rinn, J.L. Basset: learning the regulatory code of the accessible genome with deep convolutional neural networks. Genome Res 2016;26(7):990–999.

Kennedy, J. and Eberhart, R. Particle swarm optimization. IEEE Intl Conf Neural Netw 1995;4:1942–1948.

Kim, S.G., et al. EP-DNN: a deep neural network-based global enhancer prediction algorithm. Sci Rep 2016;6:38433.

LeCun, Y., Bengio, Y. and Hinton, G. Deep learning. Nature 2015;521(7553):436–444.

Liu, F., et al. PEDLA: predicting enhancers with a deep learning-based algorithmic framework. Sci Rep 2016;6:28517.

Marulanda, J.J., et al. Optimum breeding strategies using genomic selection for hybrid breeding in wheat, maize, rye, barley, rice and triticale. Theor Appl Genet 2016;129(10):1901–1913.

Meuwissen, T.H.E., Hayes, B.J. and Goddard, M.E. Prediction of total genetic value using genome-wide dense marker maps. Genetics 2001;157(4):1819–1829.

Min, S., Lee, B. and Yoon, S. Deep learning in bioinformatics. Brief Bioinforms 2017;18(5):851–869.

Poland, J. and Rutkoski, J. Advances and challenges in genomic selection for disease resistance. Annu Rev Phytopathol 2016;54:79–98.

Qiu, Z., et al. Application of machine learning-based classification to genomic selection and performance improvement. ICIC 2016;9771:412–421.

Quang, D., Chen, Y. and Xie, X. DANN: a deep learning approach for annotating the pathogenicity of genetic variants. Bioinformatics 2015;31(5):761–763.

Quang, D. and Xie, X. DanQ: a hybrid convolutional and recurrent deep neural network for quantifying the function of DNA sequences. Nucleic Acids Res 2016;44(11):e107.

Resende, M.F., Jr., et al. Accuracy of genomic selection methods in a standard data set of loblolly pine (*Pinus taeda* L.). Genetics 2012;190(4):1503–1510.

Riedelsheimer, C., Technow, F. and Melchinger, A.E. Comparison of whole-genome prediction models for traits with contrasting genetic architecture in a diversity panel of maize inbred lines. BMC Genomics 2012;13:452.

Roorkiwal, M., et al. Genome-enabled prediction models for yield related traits in chickpea. Front Plant Sci 2016;7:1666.

Rumelhart, D.E., Hinton, G.E. and Williams, R.J. Learning representations by back-propagating errors. Nature 1986;323(6088):533–536.

Schmidt, M., et al. Prediction of malting quality traits in barley based on genome-wide marker data to assess the potential of genomic selection. Theor Appl Genet 2016;129(2):203–213.

Singh, R., et al. DeepChrome: deep-learning for predicting gene expression from histone modifications. Bioinformatics 2016;32(17):i639–i648.

Spindel, J., et al. Genomic selection and association mapping in rice (*Oryza sativa*): effect of trait genetic architecture, training population composition, marker number and statistical model on accuracy of rice genomic selection in elite, tropical rice breeding lines. PLoS Genet 2015;11(2): e1004982.

Srivastava, N., et al. Dropout: a simple way to prevent neural networks from overfitting. J Mach Learn Res 2014;15:1929–1958.

Van Eeuwijk, F.A., et al. Detection and use of QTL for complex traits in multiple environments. Curr Opin Plant Biol 2010;13(2):193–205.

VanRaden, P.M. Efficient methods to compute genomic predictions. J Dairy Sci 2008;91(11):4414–4423.

Varshney, R.K. Exciting journey of 10 years from genomes to fields and markets: some success stories of genomics-assisted breeding in chickpea, pigeonpea and groundnut. Plant Sci 2016;242:98–107.

Wang, S., et al. Protein secondary structure prediction using deep convolutional neural fields. Sci Rep 2016;6:18962.

Whittaker, J.C., Thompson, R. and Denham, M.C. Marker-assisted selection using ridge regression. Genet Res 2000;75(2):249–252.

Wimmer, V., et al. Genome-wide prediction of traits with different genetic architecture through efficient variable selection. Genetics 2013;195(2):573–587.

Xiong, H.Y., et al. The human splicing code reveals new insights into the genetic determinants of disease. Science 2015;347(6218):1254806.

Xu, Y. and Crouch, J.H. Marker-assisted selection in plant breeding: from publications to practice. Crop Sci 2008;48(2):391.

Yu, X., et al. Genomic prediction contributing to a promising global strategy to turbocharge gene banks. Nat Plants 2016;2:16150.

Zeng, H., et al. Convolutional neural network architectures for predicting DNA–protein binding. Bioinformatics 2016;32(12):i121–i127.

Zhang, S., et al. A deep learning framework for modeling structural features of RNA-binding protein targets. Nucleic Acids Res 2016;44(4):e32.

Zhou, J. and Troyanskaya, O.G. Predicting effects of noncoding variants with deep learning-based sequence model. Nat Methods 2015;12(10):931–934.

Zou, C., Wang, P. and Xu, Y. Bulked sample analysis in genetics, genomics and crop improvement. Plant Biotechnol J 2016;14(10):1941–1955.

